# Severe COVID-19 associated variants linked to chemokine receptor gene control in monocytes and macrophages

**DOI:** 10.1101/2021.01.22.427813

**Authors:** Bernard Stikker, Grégoire Stik, Rudi W. Hendriks, Ralph Stadhouders

**Affiliations:** Department of Pulmonary Medicine, Erasmus MC, University Medical Center Rotterdam, the Netherlands; Centre for Genomic Regulation (CRG) and Institute of Science and Technology (BIST), Barcelona, Spain; Universitat Pompeu Fabra (UPF), Barcelona, Spain; Department of Cell Biology, Erasmus MC, Rotterdam, the Netherlands

**Keywords:** SARS-CoV-2, COVID-19, 3p21.31, GWAS, monocyte, macrophage, chemokine receptor, 3D genome organization, CTCF, gene regulation

## Abstract

Genome-wide association studies have identified 3p21.31 as the main risk locus for severe disease in COVID-19 patients, although underlying biological mechanisms remain elusive. We performed a comprehensive epigenomic dissection of the 3p21.31 locus, identifying a CTCF-dependent tissue-specific 3D regulatory chromatin hub that controls the activity of several tissue-homing chemokine receptor (CCR) genes in monocytes and macrophages. Risk SNPs colocalized with regulatory elements and were linked to increased expression of *CCR1*, *CCR2* and *CCR5* in monocytes and macrophages. As excessive organ infiltration of inflammatory monocytes and macrophages is a hallmark of severe COVID-19, our findings provide a rationale for the genetic association of 3p21.31 variants with elevated risk of hospitalization upon SARS-CoV-2 infection.

## Background

Coronavirus disease 2019 (COVID-19) is a potentially life-threatening respiratory disorder caused by the severe acute respiratory syndrome coronavirus 2 (SARS-CoV-2)[1]. Clinical manifestations of SARS-CoV-2 infection range from no or mild symptoms to respiratory failure. Life-threatening disease is often associated with an excessive inflammatory response to SARS-CoV-2, involving elevated systemic cytokine levels and profound organ infiltration by monocytes and macrophages[2, 3]. Besides clinical characteristics such as age and various comorbidities[4], genetic differences play a role in predisposing individuals to progress towards severe disease[5, 6]. In genome-wide association studies (GWASs), the 3p21.31 locus was strongly associated with increased risks of morbidity and mortality - in particular for younger (≤ 60 years) individuals[7]. However, it is currently still largely unclear how variants and genes in this locus affect the immune response against SARS-CoV-2 and COVID-19 disease pathophysiology.

## Results and discussion

COVID-19 GWAS meta-analyses (release 4 by the COVID-19 Host Genetics Initiative[8]) confirmed the strong association between the 3p21.31 locus and severe COVID-19, both when comparing hospitalized COVID-19 patients with healthy control subjects (**Fig.S1a**) or with non-hospitalized patients (**Fig.S1b**). We focused on the former comparison (8,638 hospitalized COVID-19 patients vs. 1,736,547 control subjects) to maximize the number of associated SNPs for downstream analysis. Regional association plots generated using the Functional Mapping and Annotation (FUMA) platform[9] revealed a region of 743 kb in high linkage-disequilibrium (LD; r^2^>0.8) with 958 independent significant (P<5e-8) GWAS SNPs (**Fig.1a**). Approximately 96% of these SNPs fall in non-coding regions adjacent to 12 known protein-coding genes (**Fig.1a**).

**Figure 1.**
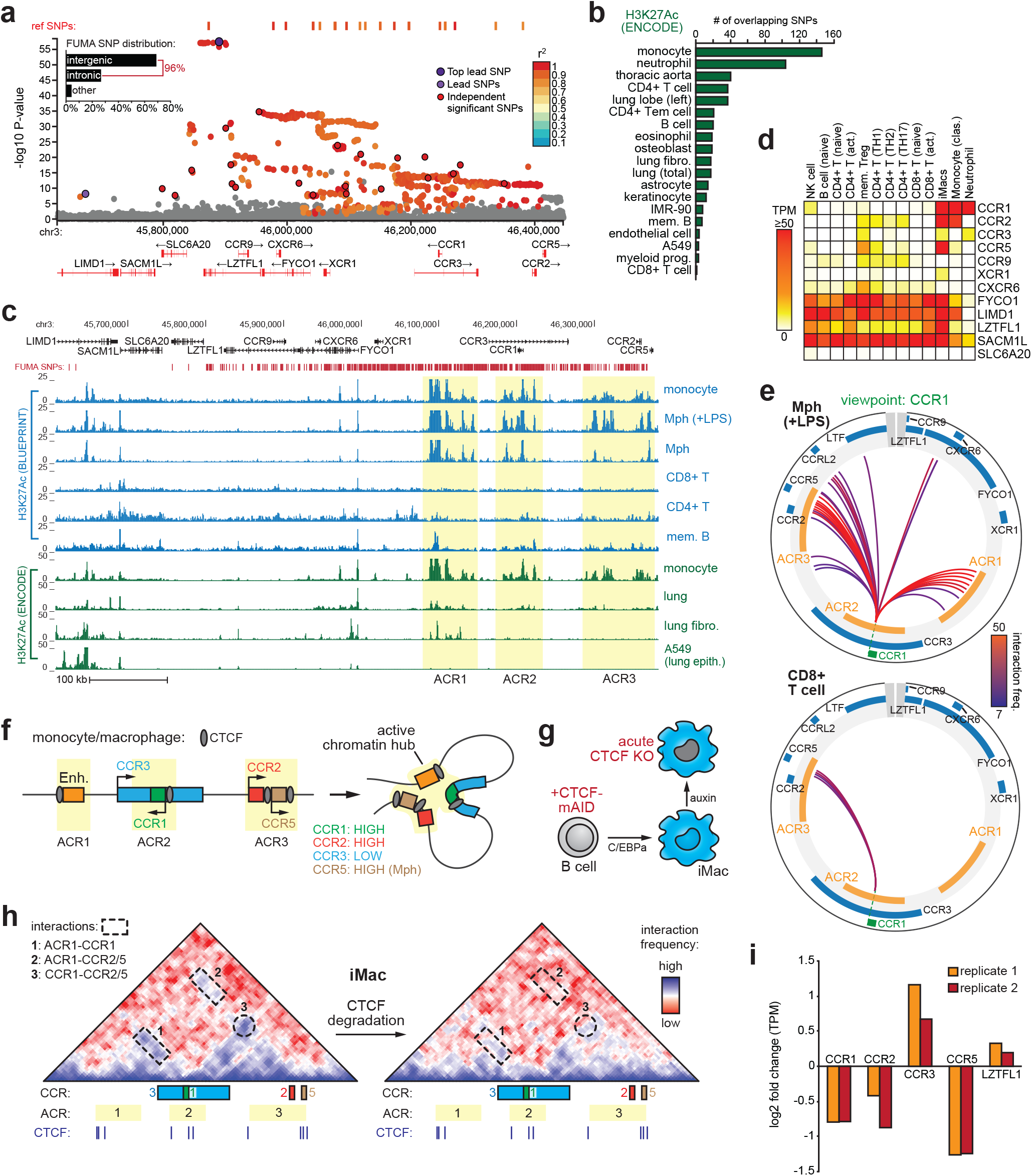
The 3p21.31 severe COVID-19 risk locus harbours a 3D chromatin hub that controls monocyte-macrophage chemokine receptor expression. (**a**) FUMA regional plot of the 3p21.31 locus highlighting all variants in high linkage-disequilibrium (r^2^>0.8, P<0.05) with independent significant (P<5e-8) GWAS SNPs. Bar graph denotes SNP distribution. (**b**) Number of COVID-19-associated SNPs overlapping with H3K27Ac^+^ regions in the indicated cell types. (**c**) UCSC genome browser view of H3K27Ac ChIP-Seq tracks for the indicated cell or tissue types (fibro. = fibroblast, epith. = epithelial, Mph = macrophage, mem. B = memory B cell). Genes and FUMA SNPs are shown above, yellow shading indicates location monocyte/macrophage-specific active chromatin regions (‘ACR1-3’). (**d**) Normalized gene expression levels (transcripts per million; TPM) of 3p21.31 candidate genes across various immune cell subsets from peripheral blood (DICE and HaemoSphere databases) and in vitro transdifferentiated induced macrophages (iMacs[16]). (**e**) Circos plots showing significant chromatin interactions with the *CCR1* promoter (green dashed line) in LPS stimulated macrophages or CD8+ T cells as measured by promoter-capture Hi-C (freq.: frequency). ACRs are indicated in orange. (**f**) Schematic indicating active chromatin hub formation involving the ACRs (enhancer; Enh.), CTCF binding sites and indicated *CCR* genes in monocytes/macrophages. (**g**) Experimental scheme depicting C/EBPα-driven transdifferentiation of B cells carrying CTCF-mAID alleles into iMacs. Exposure to auxin induces rapid degradation of CTCF-mAID[16]. (**h**) Hi-C interaction matrices (5 kb resolution, smoothened) for iMacs before (left) and after (right) auxin-inducible CTCF degradation, resulting in weaker interactions (indicated by numbers) between CCR genes and/or ACR1 (color code as in panel f). CTCF ChIP-Seq peaks in iMacs are indicated below. (**i**) Gene expression changes of indicated genes in iMacs after CTCF degradation.

Common disease-associated genetic variants predominantly localize to regulatory DNA elements[10]. To identify disease-relevant candidate genes and gene regulatory regions at 3p21.31, we integrated GWAS findings with publicly available data from large-scale transcriptomics and epigenome profiling studies. Special emphasis was placed on immune cells, as detrimental hyperinflammation is characteristic of severe COVID-19[2, 3]. Analysis of histone 3 lysine 27 acetylation (H3K27Ac) profiles from ENCODE[11] and BLUEPRINT[12] databases revealed cell type-specific active gene regulatory elements (GREs) at 3p21.31, with particularly strong activity seen in monocytes, monocyte-derived macrophages and neutrophils (**Fig.1b-c**, **Fig.S2**). The largest fraction of disease-associated SNPs overlapped with monocyte H3K27Ac^+^ GREs, which were concentrated in three active chromatin regions (ACRs) near the *CCR1*, *CCR2*, *CCR3* and *CCR5* genes (**Fig.1b-c**). CCR1 as well as CCR2 are critical mediators of monocyte/macrophage polarization and tissue infiltration[13], which are pathogenic hallmarks of severe COVID-19[2, 3]. The three ACRs also showed substantial chromatin accessibility (as measured by DNAse-Seq) in monocytes (**Fig.S2**). Gene expression analysis using data from 6 transcriptome repositories (see Methods) confirmed strong transcriptional activity of the 3’ *CCR* genes in tissues containing hematopoietic cells (e.g. whole blood, spleen), with especially *CCR1* and *CCR2* being highly expressed in classical monocytes, macrophages and neutrophils (**Fig.1d**, **Fig.S2-S3**). Of note, several other immune cell subsets, including T cell and dendritic cell subsets, also expressed specific *CCR* genes (**Fig.1d**, **Fig.S2-S3**). Chromatin interaction profiles from primary immune cells (measured by promoter-capture Hi-C[14]) revealed extensive monocyte/macrophage-specific chromatin interactions between the three ACRs, as exemplified by *CCR1* promoter interaction profiles in monocyte-derived macrophages and T cells (**Fig.1e**, **Fig.S4a-b**). In all immune cells profiled by Javierre et al.[14], no significant interactions were detected between 3p21.31 gene promoters and the lead SNP region or the most distal SNPs in *LIMD1* (**Fig.S4c**).

Together, this analysis reveals strong transcriptional activity of a *CCR* gene cluster within the 3p21.31 COVID-19 risk locus in immune cells, especially in monocytes and macrophages. Activity is centered around *CCR1* and its genomic surroundings, which are organized in a 3D chromatin hub involving the other active *CCR* genes (i.e. *CCR2*, *CCR5*) and putative enhancer elements (**Fig.1f**) - a chromatin conformation often used for complex tissue-specific gene regulation[15]. To further substantiate the relevance of local 3D chromatin organization for 3p21.31 *CCR* gene regulation in myeloid cells, we used epigenomics data from the BLaER induced macrophage (iMac) cell line system[16]. The iMacs, which morphologically and functionally closely resemble macrophages[17], showed highly comparable H3K27Ac enrichment at the 3p21.31 ACRs and expressed high levels of *CCR1*, *CCR2* and *CCR5* (**Fig.S5a-b**). High resolution in-situ Hi-C data[16] of iMacs revealed that the 3p21.31 COVID-19-associated genomic block resides in the nuclear A compartment (**Fig.S5c**), a chromosomal compartment located in the nuclear interior that groups together transcriptionally active chromatin[18]. Zooming in, we observed that most of the 3p21.31 risk variants and all associated chemokine receptor genes localize to a single topologically associating domain (TAD) (**Fig.S5d**), representing an insulated genomic neighbourhood that promotes establishing interactions between genes and regulatory elements inside the TAD[18]. Interestingly, ACR1 and ACR3 were flanked by strong binding sites for the genome architectural CCCTC-binding factor CTCF[19] in iMacs and primary monocytes (**Fig.S5a**). Together with the presence of additional CTCF binding sites within all three ACRs, including the CCR1 promoter region (**Fig.S5a**), these data suggest that CTCF organizes local 3D active chromatin hub formation to insulate the *CCR3*-*CCR1*-*CCR2*-*CCR5* gene cluster for transcriptional regulation. To test this hypothesis, we leveraged our recently developed iMac line expressing CTCF fused to an auxin-inducible degron (mAID), which allows for rapid degradation of CTCF and disruption of 3D genome architecture (**Fig.1g**)[16]. Detailed Hi-C analysis confirmed the presence of strong interactions between the ACRs and 3’ *CCR* genes in iMacs, which were disrupted upon CTCF depletion (**Fig.1h**). Importantly, chromatin hub decommissioning specifically reduced *CCR1*, *CCR2* and *CCR5* expression (**Fig.1i**), revealing that CTCF-mediated 3D chromatin interactions are critical for regulating 3p21.31 *CCR* gene activity in macrophages. Of note, expression of the *CCRL2* gene just downstream of *CCR5* - encoding an atypical chemokine receptor involved in macrophage polarization[20] - was only marginally affected by CTCF depletion (log_2_ fold change of 0.23).

We next sought to directly link COVID-19-associated genetic variants to altered 3’ *CCR* gene expression in myeloid immune cells. To this end, we used FUMA to systematically analyze previously reported expression quantitative trait loci (eQTLs) overlapping with the 958 COVID-19-associated (P<5e-8) SNPs. As eQTL sources, we focused on disease-relevant tissues rich in monocytes/macrophages (i.e. whole blood and lung tissue) and studies using purified monocytes or in vitro differentiated macrophages (see Methods). The 3’ 3p21.31 *CCR* genes showed highly significant eQTL associations (FDR<0.05) with COVID-19-associated variants, especially in monocytes and macrophages (**Fig.2a-b**). Multiple risk SNPs were identified as eQTLs for *CCR1*, *CCR2*, *CCR3* and *CCR5* in monocytes/macrophages, with the majority correlating with increased gene expression (**Fig.2c-d**). No eQTL associations were detected for *CCRL2*. To further prioritize variants with potential biological significance we used RegulomeDB[21] and CADD[22] SNP annotations. Stringent filters for both scores were combined with localization within a putative monocyte regulatory region (H3K27Ac^+^ and DNAse^+^), yielding four unique candidate causal SNPs of which three were associated with increased *CCR1*, *CCR2* and/or *CCR5* expression (**Fig.2e-f**). These variants mostly clustered within ACR2 and altered putative transcription factor binding motifs, readily providing testable hypotheses for future investigations (**Fig.S6**). For example, two SNPs within the *CCR1* promoter affected binding motifs of known regulators of the macrophage inflammatory expression program (**Fig.S6a-b**). Variant rs3181080 optimizes a composite Interferon Regulatory Factor (IRF)-Activator Protein 1 (AP1) motif, which is used for cooperative binding of IRF and AP1 family transcription factors that promote monocyte/macrophage activation[23]. In line with *CCR1* activation by IRF/AP1 factors, binding of AP1 proteins and IRF4 to rs3181080 was detected in *CCR1*-expressing GM12878 lymphoblastoid cells (**Fig.S6c**). Previous experiments in mouse macrophages[24] confirmed IRF binding to the *Ccr1* promoter (**Fig.S6d**). The second *CCR1* promoter variant, rs34919616, disrupts a critical nucleotide in a motif for BCL6 (**Fig.S6a-b**), a suppressor of inflammatory gene expression in macrophages[25].

**Figure 2.**
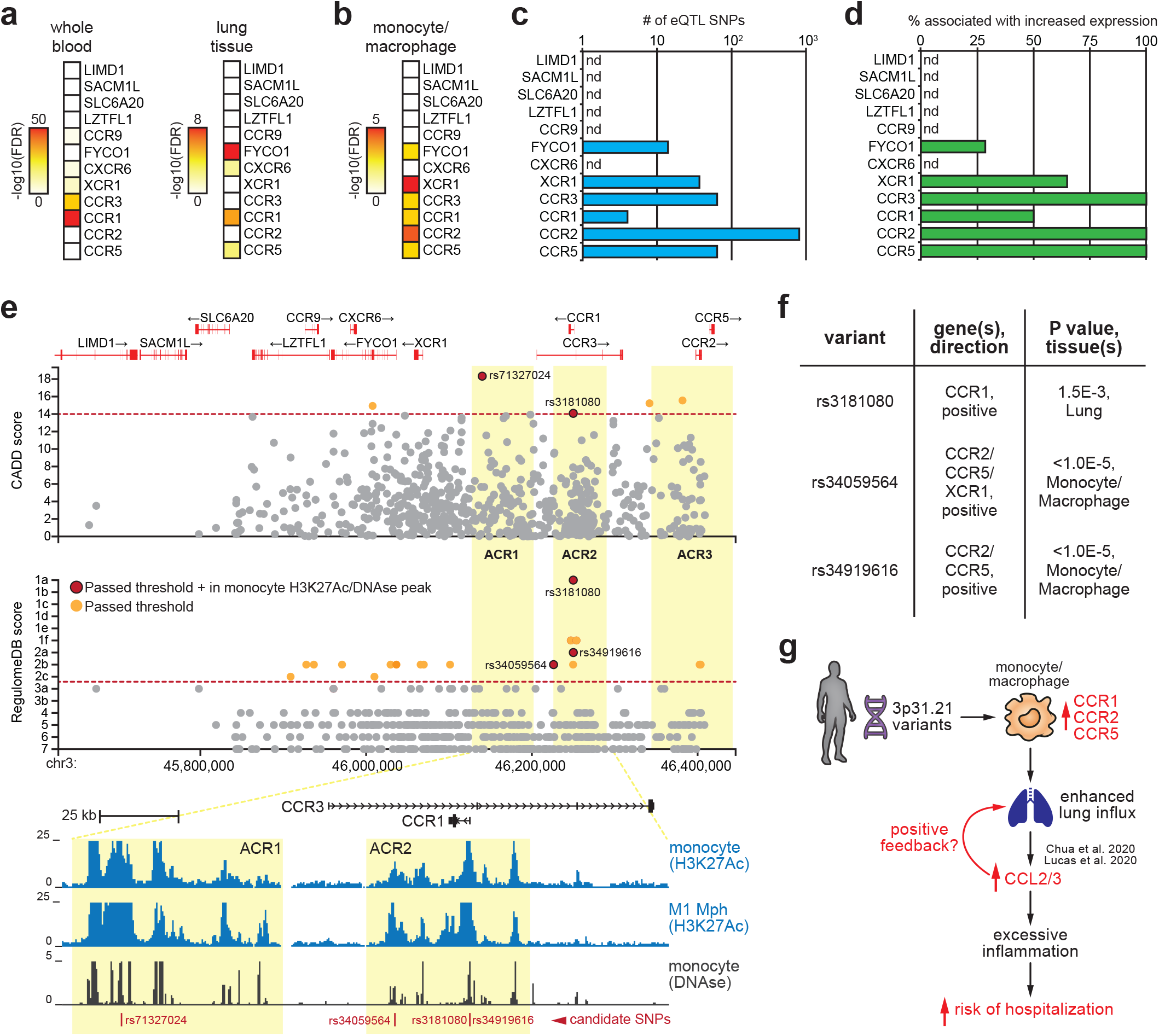
3p21.31 COVID-19 risk variants are linked to increased monocyte-macrophage chemokine receptor gene expression. (**a,b**) Heatmap depicting statistical association strength (using adjusted P-values) for the peak eQTL SNP in whole blood and lung tissue (panel a, GTEx v8) or in monocytes and macrophages (panel b, see Methods for eQTL sources). Only variants in high linkage-disequilibrium (r^2^>0.8, P<0.05) with independent significant (P<5e-8) GWAS SNPs were considered. (**c**) Number of eQTL SNPs associated with the indicated 3p21.31 genes in monocytes and macrophages. (**d**) Percentage of eQTL SNPs associated with increased expression of the indicated 3p21.31 genes in monocytes and macrophages. (**e**) CADD and RegulomeDB scores for all FUMA SNPs across the 3p21.31 locus. Red dashed lines indicate selected thresholds. SNPs passing the threshold are indicated in orange; those within monocyte H3K27Ac+/DNAse+ regulatory regions are in red. Below is depicted a UCSC genome browser view of H3K27Ac ChIP-Seq and DNAse-seq signals in the indicated cell types centered on the four candidate causal variants in ACR1 and ACR2. (**f**) eQTL analysis showing the three candidate causal variants that are also eQTLs for CCR1, CCR2 and/ CCR5. Direction of the association, P values and tissue/cell types are indicated. (**g**) Schematic indicating how 3p21.31 variants may increase risk of severe disease upon SARS-CoV2 infection through altered monocyte-macrophage chemotactic receptor expression. See text for details.

Taken together, these data show that the COVID-19-associated 3p21.31 locus harbours a CTCF-dependent tissue-specific 3D chromatin hub that controls chemotactic receptor expression in monocytes and macrophages. Several 3p21.31 variants localize to gene regulatory elements within this chromatin hub and are associated with elevated *CCR1, CCR2, CCR3* and *CCR5* expression. Mechanistically, these risk variants may modulate transcription factor binding at *CCR* gene regulatory elements. CCR1, CCR2 and CCR5 upregulation could enhance lung infiltration by monocytes and macrophages upon viral infection[13], contributing to the rapid and deleterious hyperinflammation observed in COVID-19 patients suffering from severe disease[2, 3] (**Fig.2g**). In support of this notion, single cell transcriptomics revealed increased levels of *CCR1* and *CCR5* as well as their ligands *CCL2*/*CCL3* specifically in pulmonary macrophages from critical COVID-19 patients[26, 27]. Additionally, CCL2 plasma levels showed the highest predictive value for mortality in a COVID-19 patient cohort[28]. These findings are in line with excessive pulmonary influx of monocytes and subsequent differentiation into inflammatory tissue macrophages as a hallmark of severe COVID-19 (**Fig.2g**)[26].

Our analysis has several limitations. Although we provide compelling evidence for monocyte-macrophage 3’ *CCR* gene activity linked to 3p21.31 risk variants, several other immune cell types involved in antiviral immunity also express some of these chemokine receptors (e.g. *CCR1* on neutrophils, *CCR5* on T cell subsets) and may therefore also be affected by the genetic variants. Moreover, although our analysis detected fewer non-coding regulatory activity in the 5’ part of the 3p21.31 COVID-19-associated genomic block, this region harbours the lead SNP and several actively transcribed genes with more housekeeping-like expression patterns, which may also be relevant for COVID-19 pathophysiology. Indeed, integrating loss-of-function studies in an airway epithelial carcinoma cell line with eQTL data implicated *SLC6A20* and *CXCR6* in COVID-19 pathophysiology, possibly through pleiotropic effects in multiple cell types[29]. Additionally, genetic deletion of a 68kb region around the lead SNP resulted in reduced *CCR9* and *SLC6A20* expression in a leukemic T cell line[30]. Although the lead SNPs (located near *LZTFL1*) were also reported as eQTLs for *CCR2* and *CCR5* in monocytes and macrophages, this likely reflects the high LD (r^2^>0.8) of these variants with the 3’ *CCR* SNPs (**Fig.1a**). In support of this notion, removal of the lead SNP region in a myeloid cell line did not affect 3’ 3p21.31 *CCR* gene expression[30]. Future investigations including additional (non-immune) cell types are required to identify other candidate causal genes operating in different cell types and/or under different microenvironmental circumstances.

## Conclusions

Here, we provide first evidence on how common 3p21.31 genetic variants may increase susceptibility to develop severe COVID-19: by affecting gene regulatory control of monocyte-macrophage chemotactic receptor expression. As a consequence, elevated migratory capacity of monocytes and macrophages could contribute to aggravated inflammatory responses and more severe disease. These data add to our understanding of the genetic basis of COVID-19 disease heterogeneity and support exploring therapeutic targeting of monocyte-macrophage 3p21.31 CCR activity in hospitalized COVID-19 patients.

## Methods

### GWAS data retrieval

Version 4 COVID-19 GWAS meta-analysis data was retrieved from The COVID-19 Host Genetics Initiative at https://www.covid19hg.org/. GWAS data (GRCh37/hg38 genome build) was obtained from two studies: B1_ALL (Hospitalized COVID-19 vs. non-hospitalized COVID-19; 2430 cases versus 8478 controls) and B2_ALL (Hospitalized COVID-19 vs. population; 8638 cases versus 1736547 controls). GWAS summary statistics files were used to generate input files for FUMA using standard data frame processing functions in Rstudio v.1.3.

### Identification of a high LD block of COVID-19 associated SNPs

FUMA[9] was performed for both B1_ALL and B2_ALL GWASs (version 4 summary statistics downloaded from https://www.covid19hg.org/) using default settings, with exception of the r^2^ (LD) used to define independent significant SNPs, which was set to ≥0.8. Manhattan and regional plots were generated by FUMA’s SNP2GENE function. Significant FUMA SNPs were converted to GRCh38/hg38 using UCSC LiftOver (https://genome.ucsc.edu/cgi-bin/hgLiftOver) to allow aligning variants to the epigenomic profiles.

### ChIP-Seq, DNAse-Seq and (promoter-capture) Hi-C data analysis

ChIP-Seq and DNAse-Seq epigenomic data used were retrieved from public ENCODE[11] and BLUEPRINT[12] databases. Data were visualized in the UCSC Genome Browser (https://genome.ucsc.edu). The *intersect* function of BEDTools[31] was used to determine the number of FUMA SNPs overlapping with H3K27Ac^+^ regions in the indicated cell types. Peak calling files for each H3K27Ac dataset were directly obtained from the ENCODE website (https://www.encodeproject.org/). Circos plots visualizing promoter-capture HiC data from the BLUEPRINT consortium[14] were generated using https://www.chicp.org/chicp/, with a threshold normalized interaction value of 7. ChIP-Seq and in-situ Hi-C data from *in vitro* transdifferentiated macrophages (induced macrophages or iMacs), both prior to and after auxin-inducible CTCF degradation, were obtained from GSE140528 and analyzed as previously described[16].

### Gene expression analysis

RNA-Seq profiles from a broad spectrum of selected relevant cell types were obtained from public ENCODE[11] and BLUEPRINT[12] databases and visualized in the UCSC Genome Browser. Expression value heatmaps from various collection of (immune) cell types were obtained from DICE[32] (https://dice-database.org/), GTEx v8[33] (via FUMA’s GENE2FUNCTION function), BioGPS[34] (http://biogps.org/), Haemosphere[35] (https://www.haemosphere.org/) and Monaco et al.[36] (GSE107011). Transcripts per million (TPM) values were visualized as averaged values using Morpheus (https://software.broadinstitute.org/morpheus/). RNA-Seq and TPM values for iMacs were obtained from GSE140528 and analyzed as previously described[16].

### Candidate causal variant filtering

The Combined Annotation Dependent Depletion (CADD[22]) and RegulomeDB[21] scores for all significantly associated SNPs were also obtained from FUMA. As thresholds to identify candidate causal variants, we used CADD scores >14 and RegulomeDB scores <3. SNPs were further filtered based on their combined overlap with H3K27Ac ChIP-Seq and DNAse-seq peaks in monocytes (data obtained from ENCODE[11]). Transcription factor binding motifs were obtained using HOMER[37].

### Expression quantitative trait locus (eQTL) analysis

eQTL analysis was performed using FUMA, focusing on tissues relevant for COVID-19 pathophysiology and enriched for monocytes/macrophages (i.e. whole blood and lung from GTEx v8[33]) or studies using monocytes and/or *in vitro* differentiated macrophages[38–40]. Thresholds for statistical significance were set to FDR<0.05.

## Declarations

### Ethics approval and consent to participate

Not applicable.

### Consent for publication

Not applicable.

### Availability of data and materials

All datasets generated and/or analysed during the current study are available in the public repositories and/or using the persistent web links described in the Methods section.

### Authors’ contributions

Study conception and design: B.S., R.W.H. and R.S.; Data acquisition: B.S., G.S. and R.S.; Analysis and interpretation of data: B.S., G.S., R.W.H. and R.S.; Drafting of manuscript: B.S., R.W.H. and R.S.; Critical revision: all authors.

## Acknowledgements

We thank staff of the Erasmus MC Pulmonary Medicine department for fruitful discussions. We are indebted to the COVID-19 Host Genetics Initiative for immediately making their GWAS analyses publically available, and to the genomics consortia that provide the scientific community with invaluable data.

## Figure Legends

**Figure S1. Overview of genome-wide genetic associations with severe COVID-19.** (**a,b**) Manhattan plots showing the strong association of the 3p21.31 locus with hospitalized COVID-19 (based on v4 GWAS meta-analysis release). Panel a compares hospitalized COVID-19 patients with healthy control subjects; panel b compares hospitalized with non-hospitalized patients.

**Figure S2. Transcriptional and epigenomic activity at 3p21.31 in selected cell types.** UCSC genome browser overview of RNA-Seq, DNAse-Seq and H3K27Ac ChIP-Seq tracks for the indicated cell or tissue types (BEC: bronchial epithelial cell, fibro.: fibroblast, epith.: epithelial). Yellow shading indicates location of monocyte/macrophage-specific active chromatin regions (‘ACR1-3’).

**Figure S3. Gene expression analysis of 3p21.31 candidate genes across various non-immune and immune cells types.** (**a**) Normalized gene expression levels (transcripts per million; TPM) across 54 tissues (GTEx v8 database). (**b**) Normalized gene expression levels (microarray-derived) across 84 tissues and cell types (BioGPS database). (**c**) Normalized gene expression levels (TPM) across 11 immune cell types (HaemoSphere). (**d**) Normalized gene expression levels (TPM) across 24 immune cell types (Monaco et al.[36]).

**Figure S4. Regulatory chromatin interactions across the 3p21.31 COVID-19 risk locus.** (**a**) Circos plots showing significant chromatin interactions with the CCR1 promoter (green dashed line) in monocytes, neutrophils, CD4+ T cells and total B cells measured by promoter-capture Hi-C[14] (freq.: frequency). ACRs are indicated in orange. (**b**) Significant chromatin interactions with the CCR5 promoter (green dashed line) in LPS stimulated macrophages (Mph+LPS) or CD4+ T cells. (**c**) Lack of chromatin interactions emanating from the location of the lead GWAS SNP (left circos plot) or most distal SNP in the LIMD1 gene (right circos plot). Data shown is for LPS stimulated macrophages (Mph+LPS, but identical results were obtained for the indicated cell types.

**Figure S5. Epigenomic landscape and 3D genome folding at the 3p21.31 COVID-19 risk locus in iMacs.** (**a**) UCSC genome browser view of H3K27Ac and CTCF ChIP-Seq signal for the indicated cell types. Yellow shading indicates location of monocyte/macrophage-specific active chromatin regions (‘ACR1-3’). (**b**) Normalized gene expression levels (transcripts per million; TPM) for indicated 3p21.31 genes in iMacs. (**c**) Overview of A/B compartmentalization (derived from in-situ Hi-C) surrounding the 3p21.31 severe COVID-19 associated region (indicated by orange shading) in iMacs. PC1 values (at 100kb bin size) are shown as continuous profiles, with positive values (A compartment) in green and negative values (B compartment) in blue. (**d**) In-situ Hi-C contact map (at 5kb bin size) of the 3p21.31 COVID-19 risk locus in iMacs. TAD borders called are depicted as gray rectangles (called at 50kb resolution) and the topologically associating domain (TAD) encompassing most of the severe COVID-19 associated region (indicated by orange shading) is depicted by a dashed line. H3K27Ac and CTCF enrichment in iMacs is shown below.

**Figure S6. 3p21.31 COVID-19 risk variants disrupt putative transcription factor binding sites.** (**a**) Transcription factor binding motifs overlapping with four candidate causal SNPs (see Fig.2e). Single nucleotide substitutions are indicated above the motif logos. (**b**) Zoom-in view of the CCR1 promoter (top) showing H3K27Ac, chromatin accessibility (DNAse-Seq) and binding of AP1 (i.e. ATF2, BATF, CREM) or IRF4 transcription factors in the indicated cells (ENCODE database) overlapping two candidate causal SNPs. Binding motifs modified by the variants are indicated (bottom). Rs3181080 optimizes an IRF-AP1 composite motif (A>T), while rs34919616 disrupts a BCL6 motif (G>A). (**c**) UCSC genome browser view of the CCR1 gene (located inside an intron of CCR3) showing H3K27Ac ChIP-Seq and RNA-Seq data tracks for monocytes and GM12878 lymphoblastoid cells (ENCODE database). (**d**) UCSC genome browser view of the mouse Ccr1 gene. Locations of ChIP-Seq peaks for H3K27Ac and IRF8 detected in unstimulated or lipopolysaccharide (LPS) stimulated bone marrow-derived macrophages[24] are indicated by coloured rectangles.

