## Supplementary figures and images for "Severe COVID-19 associated variants linked to chemokine receptor gene control in monocytes and macrophages"

### Supplementary Figures

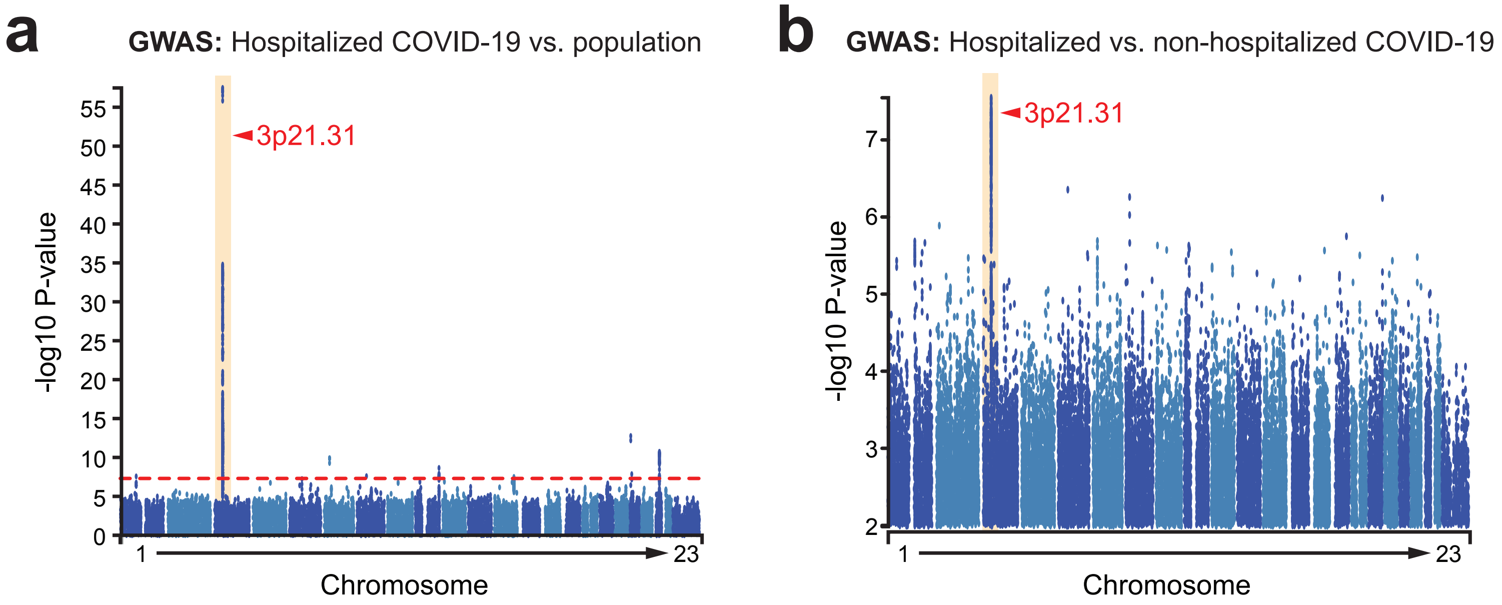

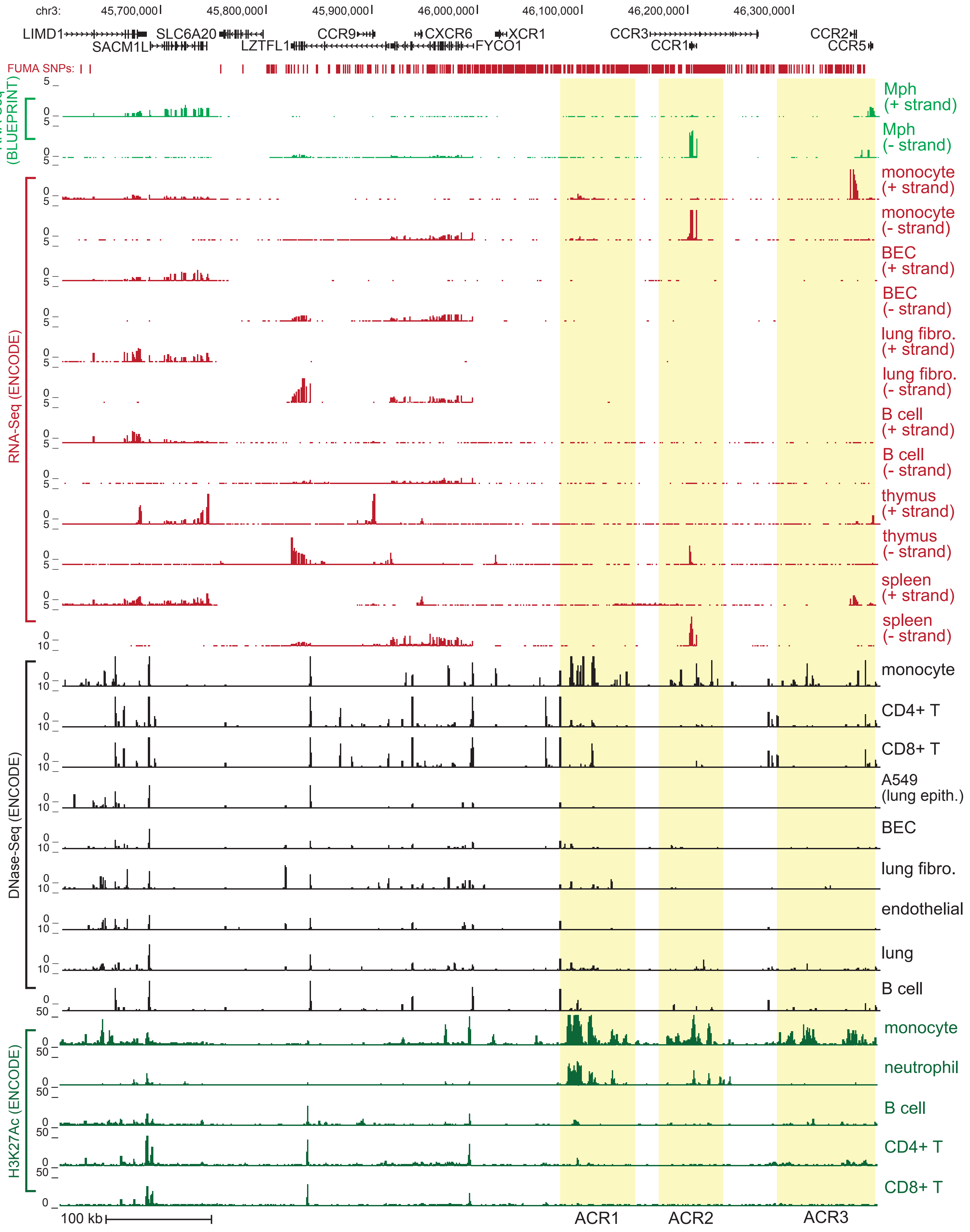

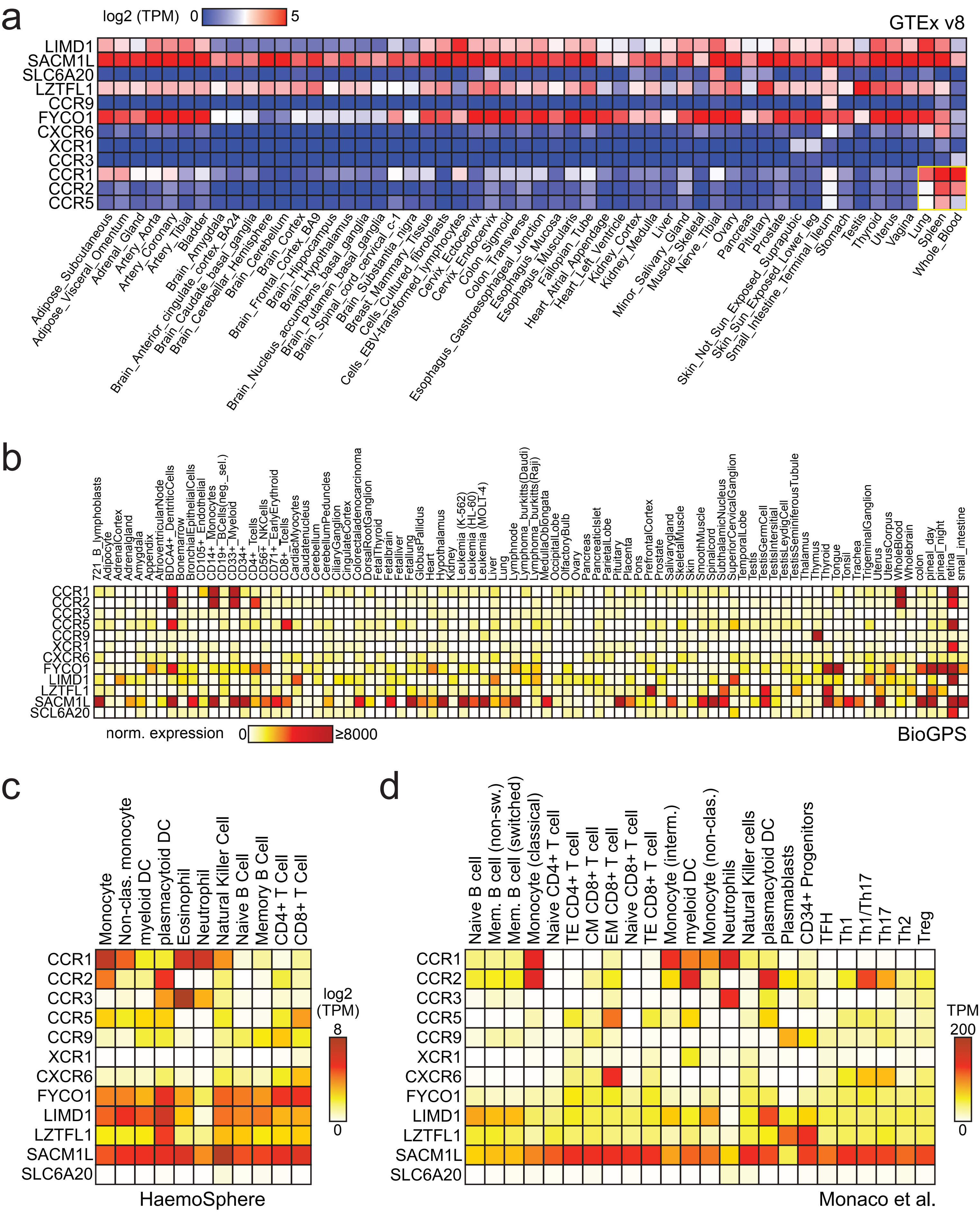

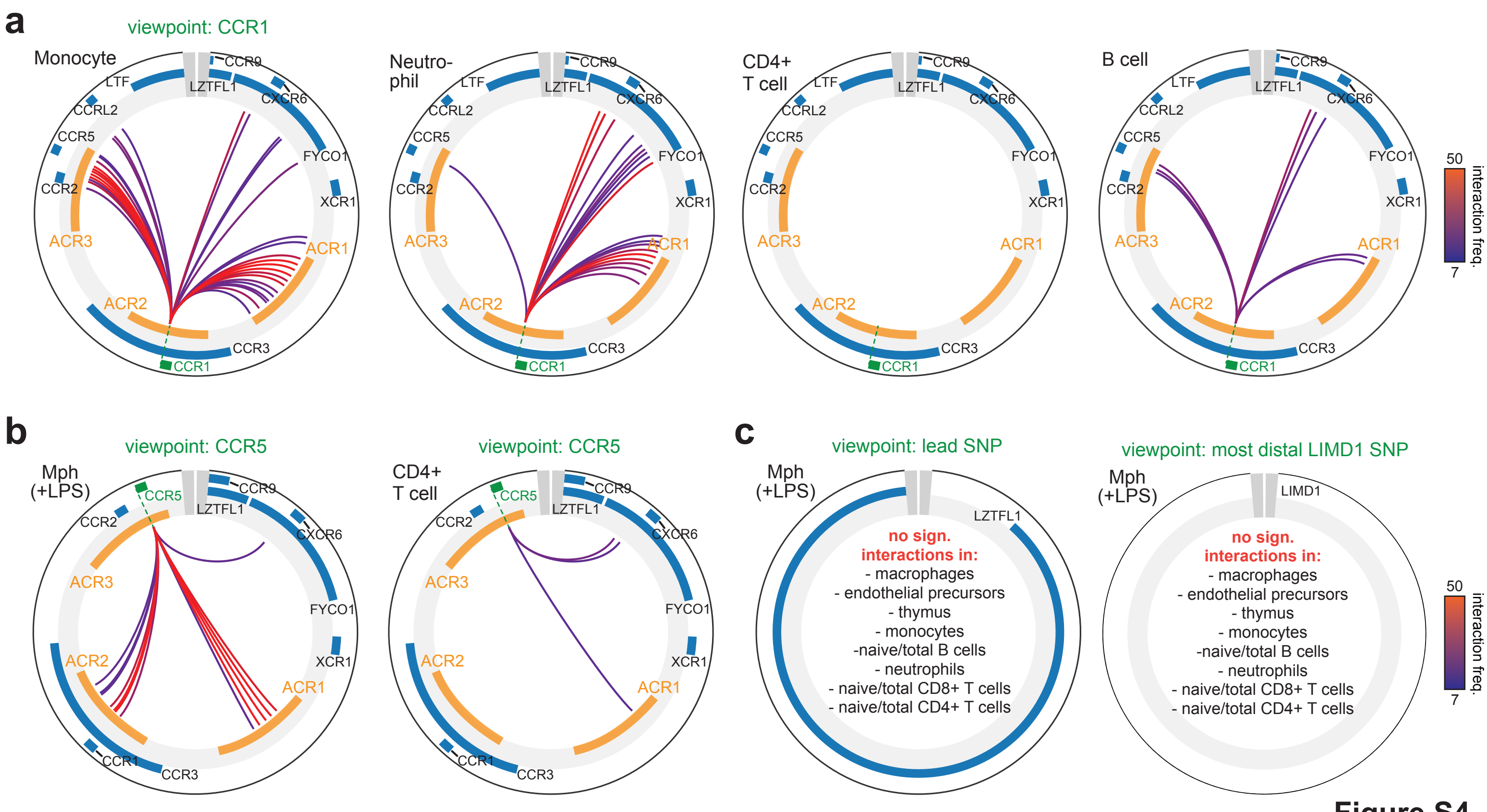

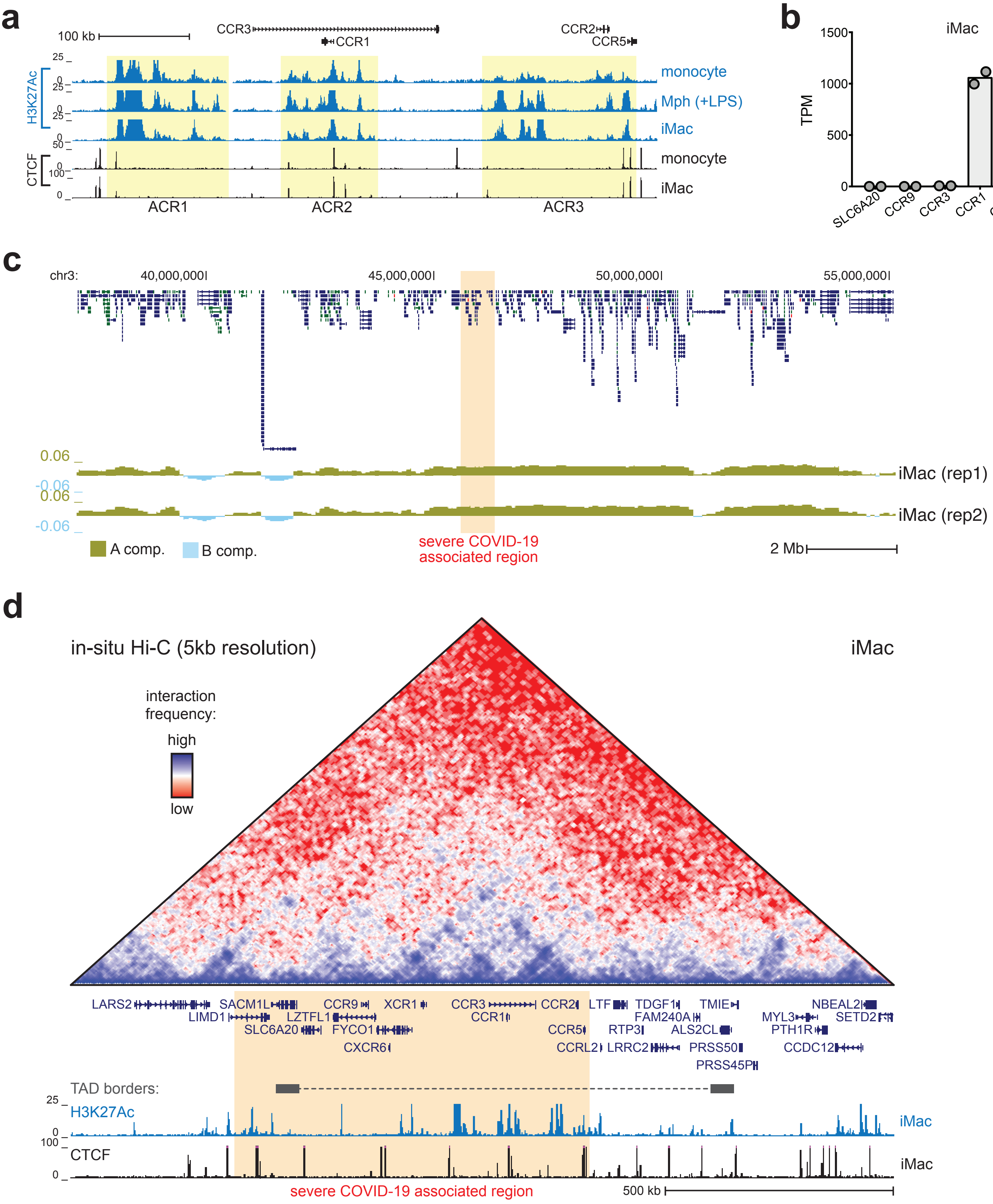

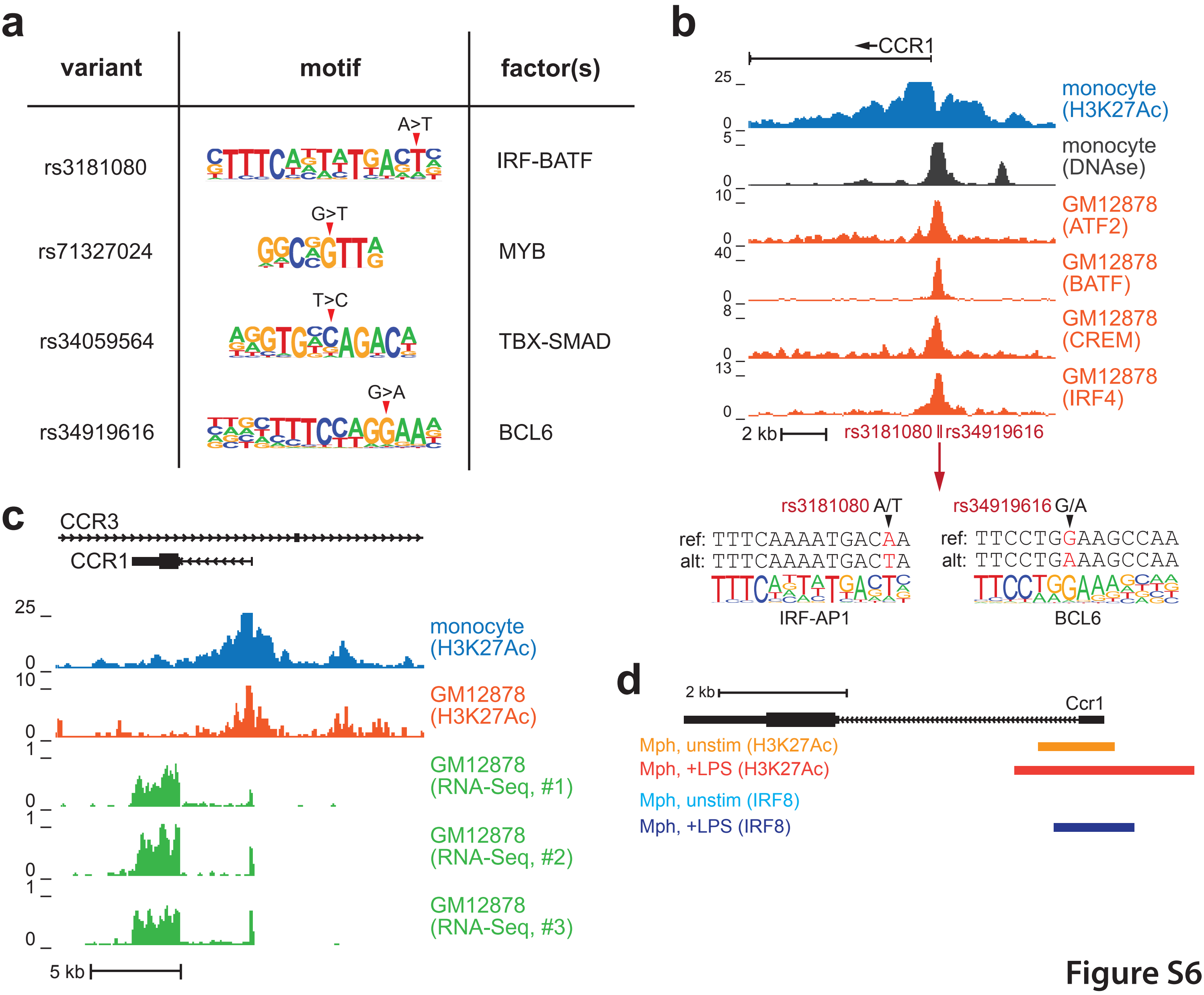
